# Using mitochondrial genomes to infer phylogenetic relationships among the oldest extant winged insects (Palaeoptera)

**DOI:** 10.1101/164459

**Authors:** Sereina Rutschmann, Ping Chen, Changfa Zhou, Michael T. Monaghan

## Abstract

Phylogenetic relationships among the basal orders of winged insects remain unclear, in particular the relationship of the Ephemeroptera (mayflies) and the Odonata (dragonflies and damselflies) with the Neoptera. Insect evolution is thought to have followed rapid divergence in the distant past and phylogenetic reconstruction may therefore be susceptible to problems of taxon sampling, choice of outgroup, marker selection, and tree reconstruction method. Here we newly sequenced three mitochondrial genomes representing the two most diverse families of the Ephemeroptera, one of which is a basal lineage of the order. We then used an additional 90 insect mitochondrial genomes to reconstruct their phylogeny using Bayesian and maximum likelihood approaches. Bayesian analysis supported a basal Odonata hypothesis, with Ephemeroptera as sister group to the remaining insects. This was only supported when using an optimized data matrix from which rogue taxa and terminals affected by long-branch attraction were removed. None of our analyses supported a basal Ephemeroptera hypothesis or Ephemeroptera + Odonata as monophyletic clade sister to other insects (i.e., the Palaeoptera hypothesis). Our newly sequenced mitochondrial genomes of *Baetis rutilocylindratus*, *Cloeon dipterum*, and *Habrophlebiodes zijinensis* had a complete set of protein coding genes and a conserved orientation except for two inverted tRNAs in *H. zijinensis.* Increased mayfly sampling, removal of problematic taxa, and a Bayesian phylogenetic framework were needed to infer phylogenetic relationships within the three ancient insect lineages of Odonata, Ephemeroptera, and Neoptera. Pruning of rogue taxa improved the number of supported nodes in all phylogenetic trees. Our results add to previous evidence for the Odonata hypothesis and indicate that the phylogenetic resolution of the basal insects can be resolved with more data and sampling effort.

## 1. Introduction

Insects are the most diverse branch of metazoan life, yet the relationships among the basal branches of the winged insects (Pterygota) remains one of the major open questions in the evolution of life. The unresolved relationship of the basal orders Odonata (dragon- and damselflies) and Ephemeroptera (mayflies) to the Neoptera (i.e., to the rest of the Pterygota) has been termed the “Palaeoptera problem” (Hovmöller, 2002; Ogden and Whiting, 2003). The name Paleoptera (“old wings”) reflects the inability of the Ephemeroptera and Odonata to fold their wings flat over the abdomen. This has long been considered to be the ancestral condition, in contrast to the more derived Neoptera (“new wings”) (reviewed by Trautwein et al. 2012). The monophyly of the Neoptera, including the three major lineages Polyneoptera, Paraneoptera, and Holometabola, is widely accepted, although relationships among polyneopteran lineages (Whitfield and Kjer, 2008) and the monophyly of the Paraneoptera (Ishiwata et al., 2011) remain unresolved.

There are three competing hypotheses relating to the Palaeoptera problem: the Palaeoptera hypothesis ((Ephemeroptera + Odonata) + Neoptera), the basal Ephemeroptera hypothesis (Ephemeroptera + (Odonata + Neoptera)), and the basal Odonata hypothesis (Odonata + (Ephemeroptera + Neoptera)) (Kjer et al., 2016; Trautwein et al., 2012; Yeates et al., 2012). All hypotheses are supported by morphological as well as molecular data to varying degrees (reviewed by Trautwein et al. 2012). Different authors, using the same set of genes and taxa set but different phylogenetic approaches, namely Bayesian inference (BI) vs. maximum likelihood (ML) (Mallatt and Giribet, 2006; Meusemann et al., 2010; Ogden and Whiting, 2003), and nucleotide vs. amino acid (aa) sequences (Song et al., 2016) obtained results supporting distinct hypotheses. The Palaeoptera hypothesis has received support from several molecular studies based on rRNA genes and nuclear DNA (Ishiwata et al., 2011; Kjer et al., 2006; Sasaki et al., 2013; Thomas et al., 2013), and from nDNA phylogenomic analyses (Meusemann et al., 2010; Regier et al., 2010; Reumont et al., 2012). Thomas et al. (2013) and Kjer et al. (2006) included the most comprehensive taxon sampling with 35, respectively seven species from each of the three lineages (i.e., Ephemeroptera, Odonata, Neoptera) analyzing seven, respectively eight genes (rRNAs, nuclear DNA (nDNA), mitochondrial DNA (mtDNA)). The basal Ephemeroptera hypothesis was supported by previous studies using mitochondrial genome data (Ma et al., 2014; Song et al., 2016; Zhang et al., 2008), and a combined analysis of rRNA and one nuclear gene (Ogden and Whiting, 2003). Other studies have concluded the relationships are ambiguous (Ogden and Whiting, 2003). Notably, studies supporting the basal Ephemeroptera hypothesis have included either a limited number of mayfly species (e.g., one (Whiting et al., 1997; Zhang et al., 2008), two (Giribet and Ribera, 2000; Ma et al., 2014), four (Ma et al., 2015), or five (Song et al., 2016) species) or a limited number of unlinked loci (e.g., one (Ma et al., 2014; Ma et al., 2015; Song et al., 2016; Zhang et al., 2008), two (Giribet and Ribera, 2000; Wheeler et al., 2001; Whiting et al., 1997), or three (Ogden and Whiting, 2003)). The basal Odonata hypothesis has received support from previous studies based on mitochondrial genome data (Lin et al., 2010; Wan et al., 2012), rRNA genes (Kjer, 2004; Mallatt and Giribet, 2006; Misof et al., 2007; Reumont et al., 2009; Yoshizawa and Johnson, 2005), combining rRNA and nDNA (Blanke et al., 2013), and phylogenomic nDNA (Meusemann et al., 2010; Simon et al., 2009). These studies, with the exceptions of few studies based on *18S* rRNA (Kjer, 2004; Misof et al., 2007; Yoshizawa and Johnson, 2005), included in total between two (Mallatt and Giribet, 2006; Simon et al., 2009) and 13 species (Reumont et al., 2009) from the orders Ephemeroptera and Odonata.

The conflicting phylogenetic signals may result from the ancient radiation of the lineages Ephemeroptera, Odonata, and Neoptera from a common ancestor in the distant past (Whitfield and Kjer, 2008). These three ancient lineages appear to have diverged rapidly, leaving few characteristics to determine their phylogenetic relationships (Whitfield and Kjer, 2008). Evolutionary rate heterogeneity across clades and the representation of old clades by recent extant taxa make ancient relationships such as those of early winged insects the most difficult challenges for phylogenetics (Kjer et al., 2016; Whitfield and Kjer, 2008). Given the weak phylogenetic signal, reconstructions are more susceptible to systematic errors (Baurain et al., 2007; Rokas and Carroll, 2006; Thomas et al., 2013; Whitfield and Kjer, 2008; Whitfield and Lockhart, 2007). Other sources of conflicting signal in studies of the relationships among these three lineages may result from differences in taxon sampling, sequence data, alignment methods, and phylogenetic methods used including models of evolution (Talavera and Vila, 2011; Thomas et al., 2013; Trautwein et al., 2012). The use of mitochondrial protein-coding genes (PCGs) derived from mitochondrial genomes can circumvent some of these difficulties, since they are easier to align and also appropriate models of molecular evolution are well established (Abascal et al., 2006; 2005; Le et al., 2017). On the other hand, mitochondrial genes include several drawbacks, most importantly the possible presence of pseudogenes (Bensasson, 2001; Rogers and Griffiths-Jones, 2012; Song et al., 2008). However, data derived from high-throughput sequencing, depending on the coverage bears the potential to distinguish between functional genes and pseudogenes based on higher mutation rates of the later (Claes and De Leeneer, 2014).

Overall, mitochondrial markers are well studied and therefore the most widely employed genetic markers in insects, being considered a promising “instrument” for insect systematics (Cameron, 2014). Insect mitochondrial genomes are highly conserved, ranging from 15 to 18 kb in length, containing 37 genes: 13 PCGs, two ribosomal RNAs (rRNAs, *rrnL* and *rrnS*), and 22 transfer RNAs (tRNAs, *trn\**) (Boore, 1999). A non-coding region of variable length, thought to be the origin of initiation of transcription and replication, is typically present (Saito, 2005; Zhang and Hewitt, 1997) and referred to as the AT-rich region. The typical ancestral insect mitochondrial genome differs from the ancestral arthropod mitochondrial genome only by the location of *trnL* (Boore et al., 1995). Significant differences in structure, gene content, and gene arrangement have been found to be the exception for highly derived taxa.

Importantly there is still an unbalance in the numbers of available molecular markers between the different insect lineages. In particular the early winged insects (i.e., Ephemeroptera and Odonata) are relatively under-represented. This is (partly) because so far most genomic data were obtained for insect orders that are of economical interest (e.g., as pollinators, model species, agricultural pests, and vectors of human diseases, Robinson et al., 2011). As of January 2017, there were 39 complete or nearly complete palaeopteran mitochondrial genomes (Ephemeroptera: 18, Odonata: 21) from 22 families (eleven of each) available on GenBank; 22 of which were included in publications (Ephemeroptera: ten, Odonata: twelve). Compared to the number of families that have been described (Ephemeroptera: 42 (Barber-James et al., 2008), Odonata: 30 (Dijkstra et al., 2013)) this is still a relatively low number.

The insect order Ephemeroptera comprises over 3,000 species, comprising as the two most species rich families the Baetidae (833 species) and the Leptophlebiidae (608 species) (Barber-James et al., 2008). Their biogeographic origin is probably Pangean, including a greater diversity and higher endemism rate in the Neotropics and Australasia for the Baetidae, and the Neotropical and Afrotropical regions for the Leptophlebiidae, respectively (Barber-James et al., 2008).

We investigated the relationships of the oldest extant winged insects and newly sequenced three mayfly mitochondrial genomes to improve sampling of this under-represented order. New taxa included one representative of the family Leptophlebiidae, for which no mitochondrial genome data were available, and two representatives of the Baetidae, one from each subfamily. We used BI and ML approaches and created matrices that eliminated taxa that reduced overall tree support (rogue taxa) and that were affected by long-branch attraction (LBA). Our analysis included 29 palaeopteran mitochondrial genomes from 20 families and 64 other insect mitochondrial genomes. We expect the increased taxon sampling to improve the phylogenetic signal of the basal nodes and add support for one of the three competing hypotheses.

## 2. Material and methods

### 2.1. Taxon sampling

We newly sequenced three mitochondrial genomes of the two most diverse Ephemeroptera families. *Habrophlebiodes zijinensis* GUI, ZHANG & WU, 1996 is the first representative of the family Leptophlebiidae to have its complete mitochondrial genome sequenced. *Baetis rutilocylindratus* WANG, QIN, CHEN & ZHOU, 2011 and *Cloeon dipterum* L. 1761 are both members of the Baetidae and representative each subfamily: *B. rutilocylindratus* from Baetinae, and *C. dipterum* from Cloeninae. There was a mitochondrial genome from one other member of the Baetinae available on GenBank prior to our study (*Alainites yixiani*, GQ502451, Jia & Zhou). Our analysis also included four species from Archaeognatha, three from Zygentoma (formerly Thysanura), ten from Odonata, 16 additional species from Ephemeroptera, four from Plecoptera, four from Ensifera of Orthoptera, nine from Caelifera of Orthoptera, 13 from Phasmatodea, two from Mantodea, six from Blattodea, 13 from Isoptera, and one each from Collembola, Dermaptera, Grylloblattodea, and Mantophasmatodea (Table 1). Amino acid sequences were obtained from GenBank using a custom Python script (mitogenome_ncbi.py, https://github.com/srutschmann/python_scripts).

**Table 1.**
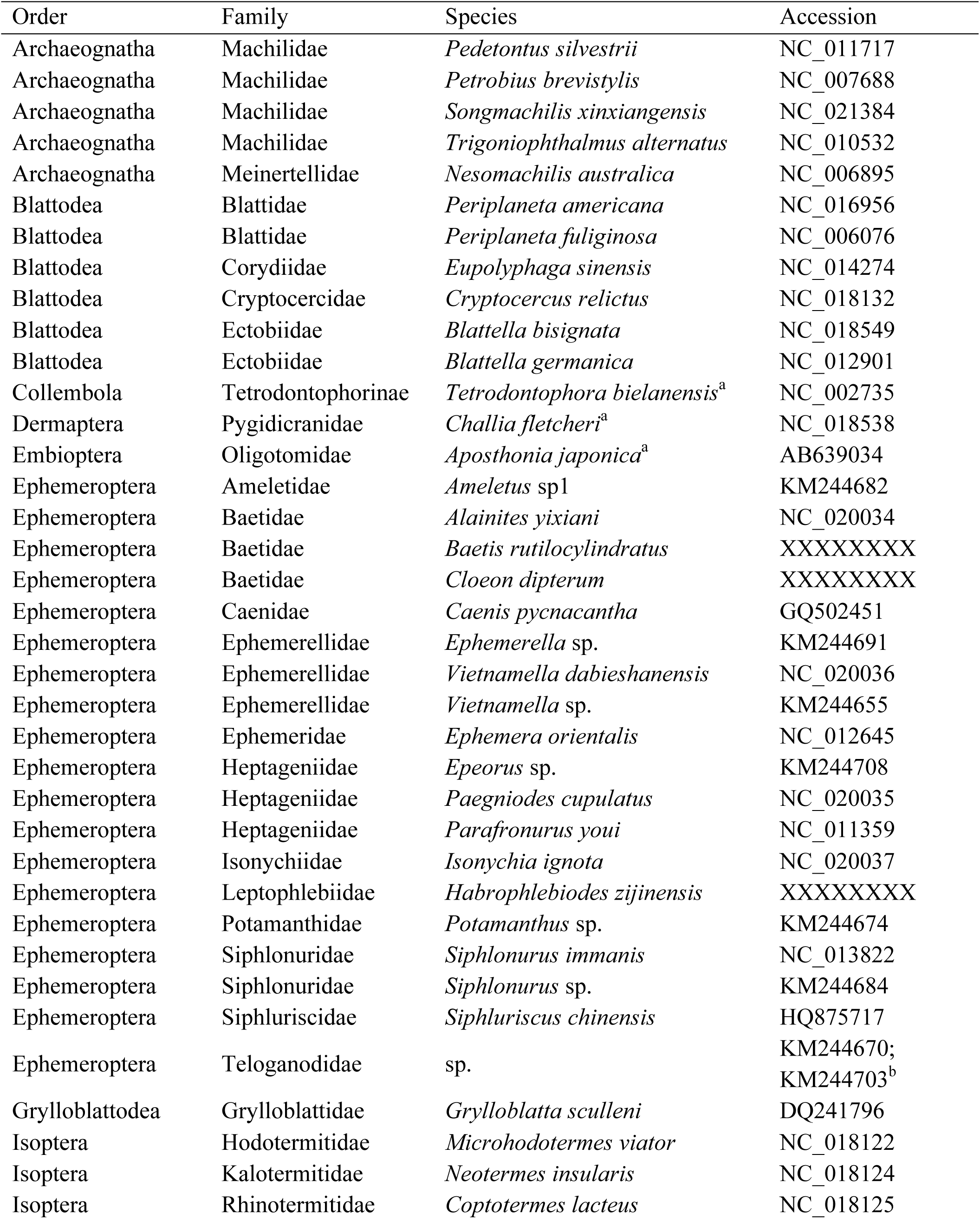
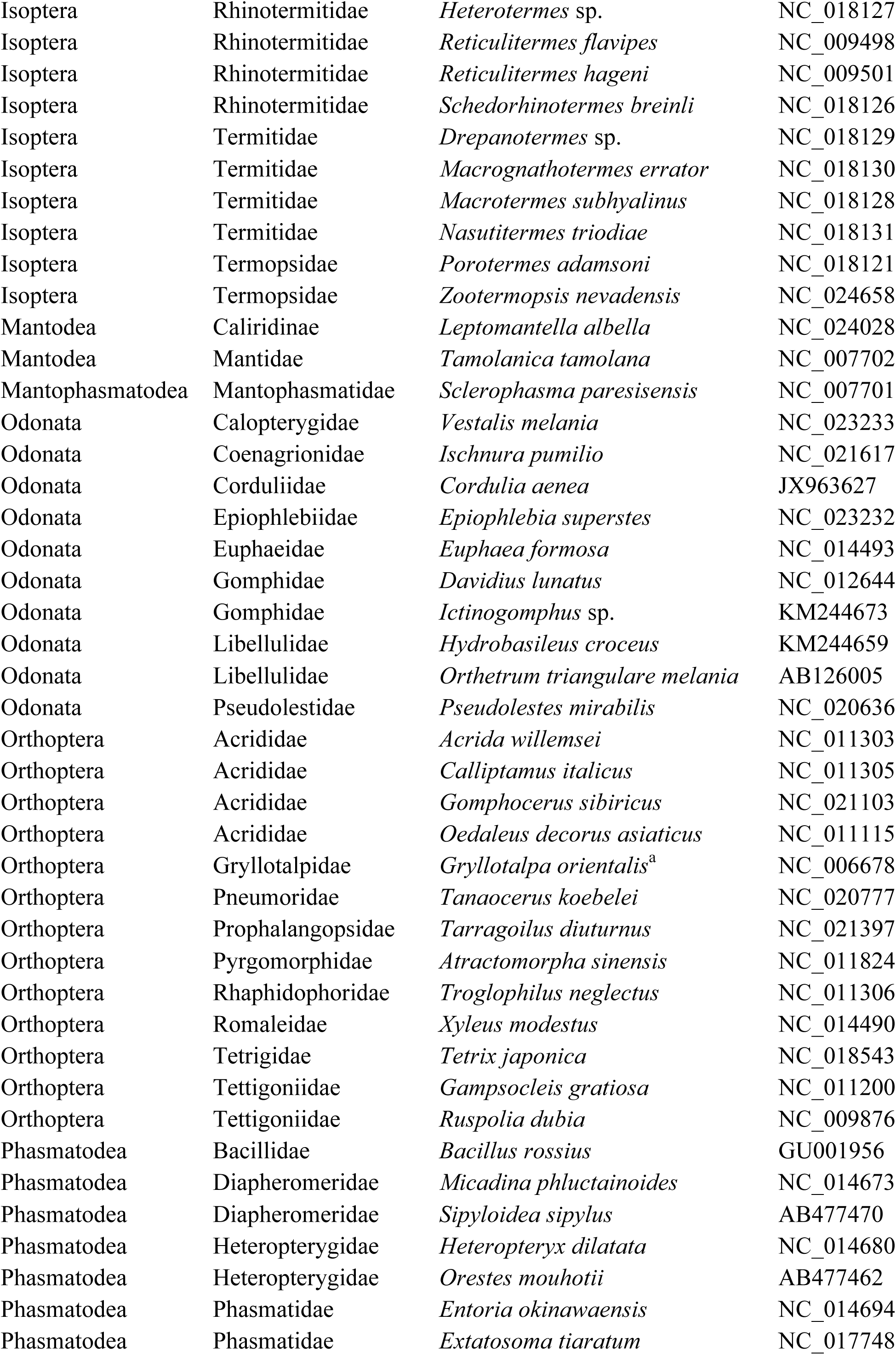
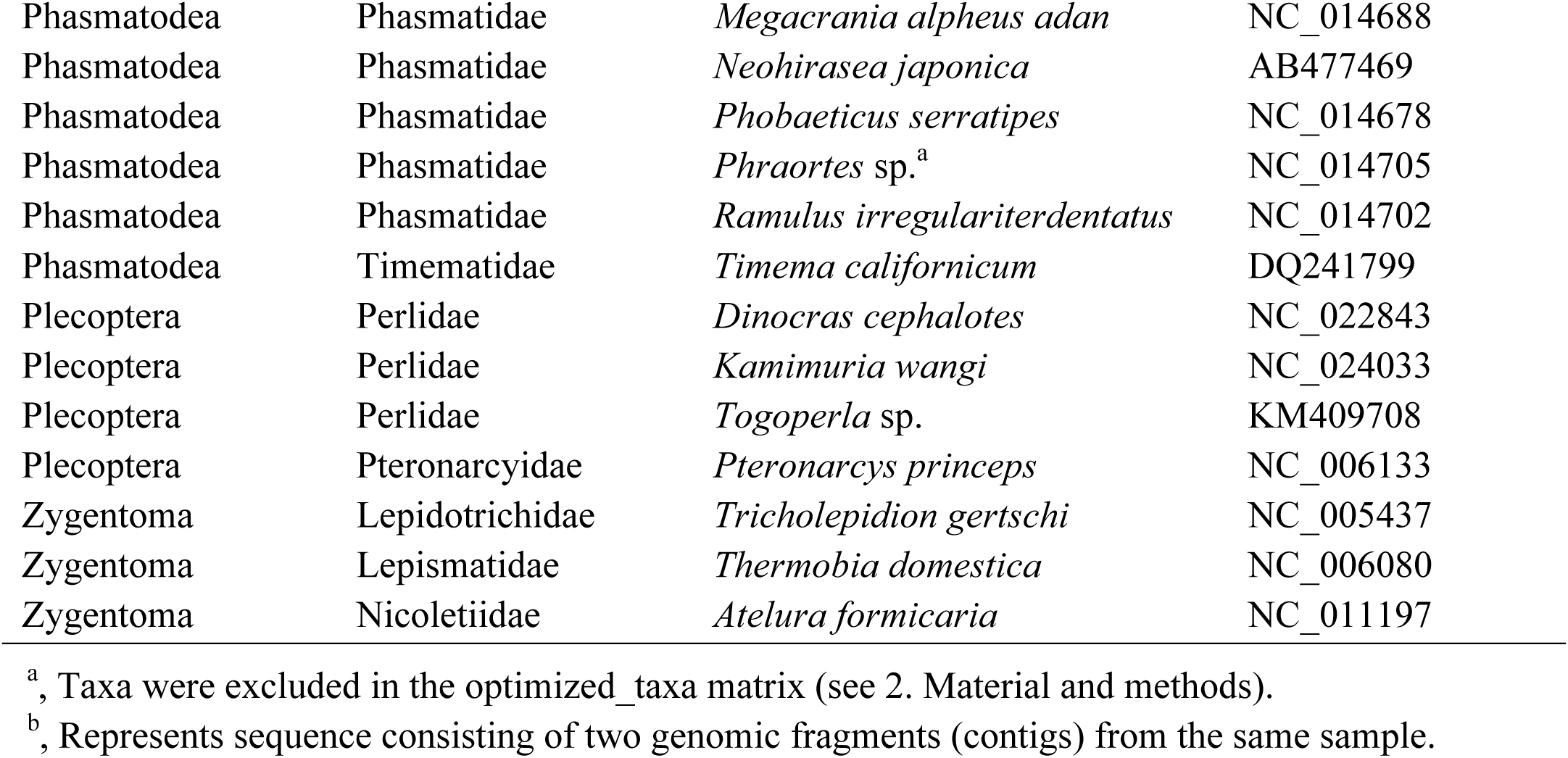
Set of mitochondrial genomes with according GenBank accession numbers.

### 2.2. Sequencing and assembly

The three mayfly species were sequenced with either 454/FLX pyrosequencing (*C. dipterum*) or Sanger sequencing (*B. rutilocylindratus* and *H. zijinensis*). We extracted DNA from *C. dipterum* from twelve to twenty pooled reared subimago specimens, prepared a shotgun library, and sequenced four lanes on a Roche (454) *GS FLX* machine at the Berlin Center for Genomics in Biodiversity Research (BeGenDiv, Berlin, Germany) (see also Rutschmann et al., 2017). The obtained sequence reads were trimmed and *de novo* assembled using the software Newbler v.2.5.3 (454 Life Science Cooperation) under default settings for large data sets. In order to extract the mitochondrial genome of *C. dipterum*, we performed BLASTN searches (Altschul, 1997) using as query all assembled contigs against the NCBI database. We mapped all matching reads back to the mitochondrial genome with BWA (Li and Durbin, 2009), using the BWA-SW algorithm (Li and Durbin, 2010) with settings suggested for 454 data by CORAL (match score = 2, mismatch penalty = 2, and gap open penalty = 3, Salmela and Schroder, 2011).

Specimens of *B. rutilocylindratus* and *H. zijinensis* were collected in Zijin Hill, Nanjing, China. The DNA was extracted from between two and four larvae using the DNeasy® Blood & Tissue (Qiagen, Leipzig, Germany) kit. Four DNA fragments of each species were amplified with universal primers (*B. rutilocylindratus*: *cob*, *cox1*, *cox3*, *rrnL*; *H. zijinensis*: *cob*, *cox1*, *nad4*, *rrnL*, (Simon et al., 1994)). Subsequently, based on the previously obtained sequence information we designed six (*B. rutilocylindratus*) respective four (*H. zijinensis*) specific primer pairs (Supplementary Table S1; see also Li et al. (2014)). Standard and long polymerase chain reactions (PCRs) were performed on a DNA Engine Peltier Thermal Cycler (Bio-Rad, Shanghai, China). Therefore, we used the rTaq^TM^ DNA polymerase (TaKaRa Bio, Dalian, China) for fragments smaller than two kb and the LA Taq^TM^ polymerase (TaKaRa Bio, Dalian, China) for fragments larger than two kb. All PCR products were purified with the Axygen agarose-out kit. When the PCR amplification signal was too weak to sequence or sequencing resulted in overlap peaks, the products were ligated to pGEM®-T Easy Vector (Promega, Southampton, UK) by *Escherichia coli*, and each resulting clone was sequenced. All purified amplification products were sequenced successively in both directions on an ABI3130xl capillary sequencer. Forward and reverse sequences were assembled and edited using CodonCode Aligner v.3.5.6 (CodonCode Corporation).

### 2.3. Sequence alignments and data matrices

We prepared two data matrices of aa sequences based on the annotated mitochondrial genomes (see 2.5. Annotation and characterization), including a set with all 93 taxa (all_taxa matrix), and a matrix for which we removed taxa that were either identified as rogue taxa using the RogueNaRok algorithm (Aberer et al., 2012; Wilkinson, 1996) and showing LBA (Felsenstein, 1978; Hedtke et al., 2006) (optimized_taxa matrix; 88 taxa, Table 1). We used GUIDANCE2 (Landan and Graur, 2008; Sela et al., 2015) to align the sequences, and identify and remove positions detected as ambiguously aligned regions. To run GUIDANCE we applied the default settings, using as multiple sequences alignment program MAFFT v.7.050b (L-INS-I algorithm with default settings, Katoh and Standley, 2013).

The best-fitting partitioning schemes and corresponding aa substitution models were estimated with PartitionFinder v.2 (https://github.com/brettc/partitionfinder, Lanfear et al., 2012). We used the Bayesian Information Criterion (BIC) to choose the best model, linked branch lengths, and a greedy search (Lanfear et al., 2012) with RAxML v.8.2.8 (Stamatakis, 2014). Thereby each gene was defined as one data block (i.e., possible partition).

To identify rogue taxa, we applied the RogueNaRok algorithm (Aberer et al., 2012; Wilkinson, 1996). We used the bootstrap replicates and the best supported maximum likelihood tree based on the all_taxa matrix (see 3.1. Data matrices). Identified rogue taxa were removed from the all_taxa matrix, resulting in the optimized_taxa matrix (Table 1). Taxa showing evidence of LBA *sensu* Bergsten (Bergsten, 2005) were identified using heterogeneities in sequence divergence based on a model of evolution derived from the entire data matrix. For this, the Relative Composition Frequency Variability (RCFV) values of each species were calculated with BaCoCa (Kück and Struck, 2014) using the partitioned concatenated data matrix (i.e., all_taxa matrix). Taxa with the highest values were excluded in the optimized_taxa matrix, apart from the Ephemeroptera, which were retained because they are the focus of our study and their monophyly is well-established (Hovmöller, 2002; Ogden and Whiting, 2003).

### 2.4. Phylogenetic reconstruction

Phylogenetic reconstructions were carried out using MrBayes v.3.2.4 (Ronquist et al., 2012), and RAxML v.8.1.7 (Stamatakis, 2014). We analyzed both matrices (all_taxa, optimized_taxa) using the best-fitting partitioning scheme and model for each (Table 2). For BI, we unlinked the frequencies, gamma distributions, substitution rates and the proportion of invariant sites across partitions. For each matrix, we run two independent analyses of four MCMC chains with 10^7^ generations, sampling every 10^3^ generations and discarding a burn-in of 25%. Maximum likelihood inferences were performed with 200 bootstrap replicates. Trees reconstructed from the all_taxa matrix were rooted with *Tetrodontophora bielanensis* (Collembola). Trees reconstructed from the optimized_taxa matrix were rooted with *Nesomachilis australica* (Archaeognatha, according to Song et al. 2016) because *Tetrodontophora bielanensis* was found to be a rogue taxon (see 3. Results). All trees were visualized with the ape v.4.1 (Paradis et al., 2004), phangorn v.2.1.1 (Schliep, 2011), and ggtree v.1.4.20 (Yu et al., 2017) packages for R v.3 (R Development Core Team, 2012).

**Table 2.**
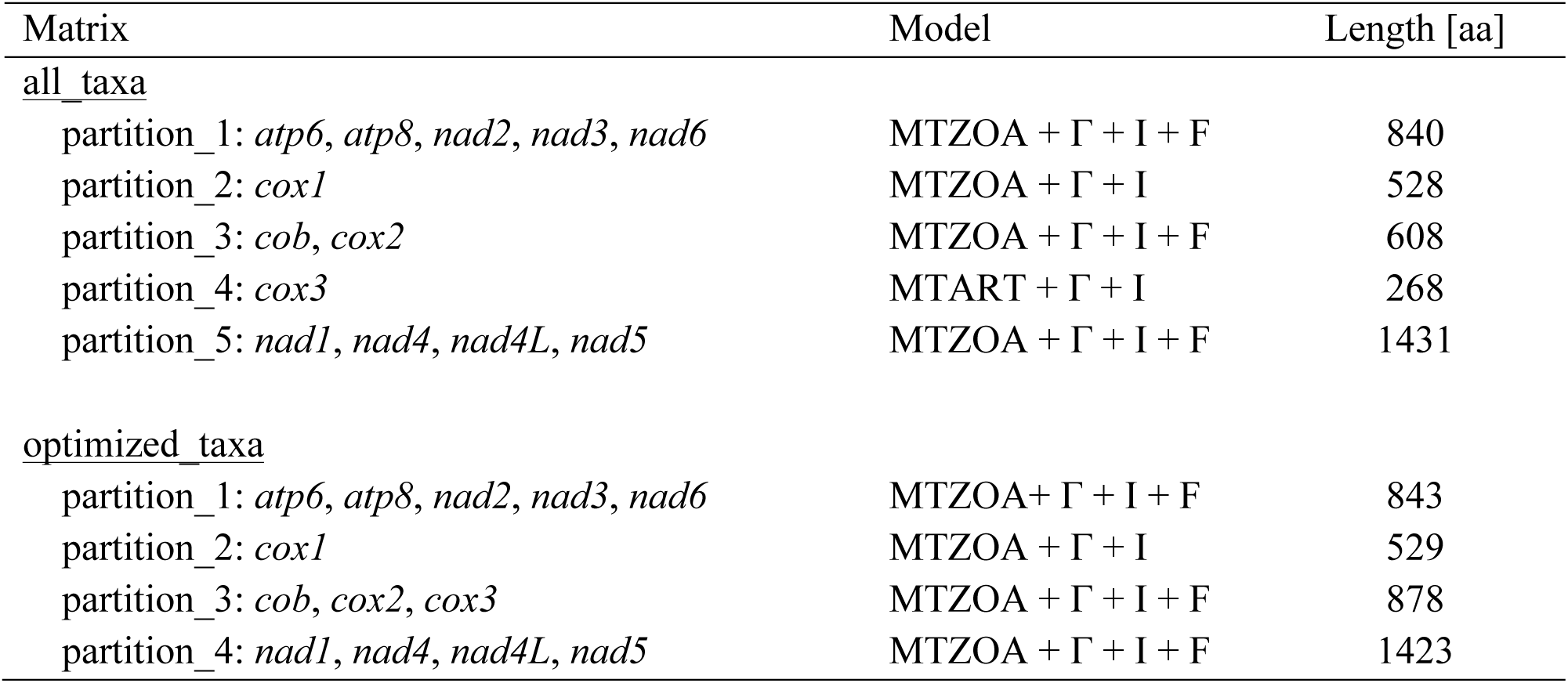
Overview of data matrices, including the best-fitting partitioning and model schemes, and amino acid (aa) sequence length.

### 2.5. Annotation and characterization

All mitochondrial genomes were annotated using the MITOS webserver (http://mitos.bioinf.unileipzig.de/index.py, Bernt et al., 2013), DOGMA (Wyman et al., 2004), and Mfannot (http://megasun.bch.umontreal.ca/cgi-bin/mfannot/mfannotInterface.pl). The predicted PCGs were checked for stop codons and manually adjusted by comparison with the longest predicted open reading frames as implemented in Geneious R7 v.7.1.3 (Biomatters Ltd.), and by comparison with the respective homologous insect sequence alignment. For the tRNA prediction, we additionally used ARWEN v.1.2.3 (Laslett and Canback, 2008) and tRNAscan-SE v.1.21 (http://lowelab.ucsc.edu/tRNAscan-SE/, Lowe and Eddy, 1997). The annotated mitochondrial genomes were visualized with OrganellarGenomeDRAW (http://ogdraw.mpimp-golm.mpg.de/cgi-bin/ogdraw.pl, Lohse et al., 2013) and manually edited.

Nucleotide contents were retrieved using Geneious. To correct biases in the AT content due to incomplete mitochondrial genomes mostly missing the AT-rich region, we removed all AT-rich regions and the two rRNAs including the five close-by tRNAs (*trnL1-trnM*) and recalculated the nucleotide base pair (bp) compositions. The AT and GC composition skewness were calculated as follows: AT-skew = (A - T) / (A + T), and GC-skew = (G - C) / (G + C) (Perna and Kocher, 1995).

## 3. Results

### 3.1. Data matrices

The final matrices had lengths of 3,675 aa (all_taxa matrix, including 93 taxa) and 3,673 aa (optimized_taxa matrix, including 88 taxa, Table 1). As best-fitting partitioning scheme for the two matrices, we identified either five partitions (all_taxa matrix, Table 2) or four partitions (optimized_taxa, Table 2). As best-fitting models of aa sequence evolution, we identified MtZoa (Rota-Stabelli et al., 2009) and MtArt (Abascal et al., 2006).

The RogueNaRok analysis identified the outgroup *T. bielanensis* as well as *Gryllotalpa orientalis* (Orthoptera) and *Phraortes* sp. (Phasmatodea) as taxa with uncertain phylogenetic positions, leading to less accurate phylogenetic reconstructions. *G. orientalis* (Orthoptera) and *Phraortes* sp. (Phasmatodea) clustered within the corresponding order using the all_taxa matrix, although the nodes containing these species were not supported by ML. In the test for LBA, *Aposthonia japonica* (Embioptera) and *Challia fletcheri* (Dermaptera) had high RCFV values (0.0237 and 0.0307, respectively) suggesting they were affected by LBA. Both species clustered within other orders: *Challia fletcheri* (Dermaptera) within the Ephemeroptera as sister taxon to a clade containing all representatives of the family Baetidae, and *A. japonica* (Embioptera) within the Phasmotodea as sister taxon to a clade containing all species of the order with the exception of *T. californicum* (Table 3, Supplementary Fig. S1). The mayflies Teloganodidae sp. (0.0305) and the three baetids *A. yixiani* (0.0235), *B. rutilocylindratus* (0.0232), and *C. dipterum* (0.0201) showed relatively high RCFV values.

**Table 3.**
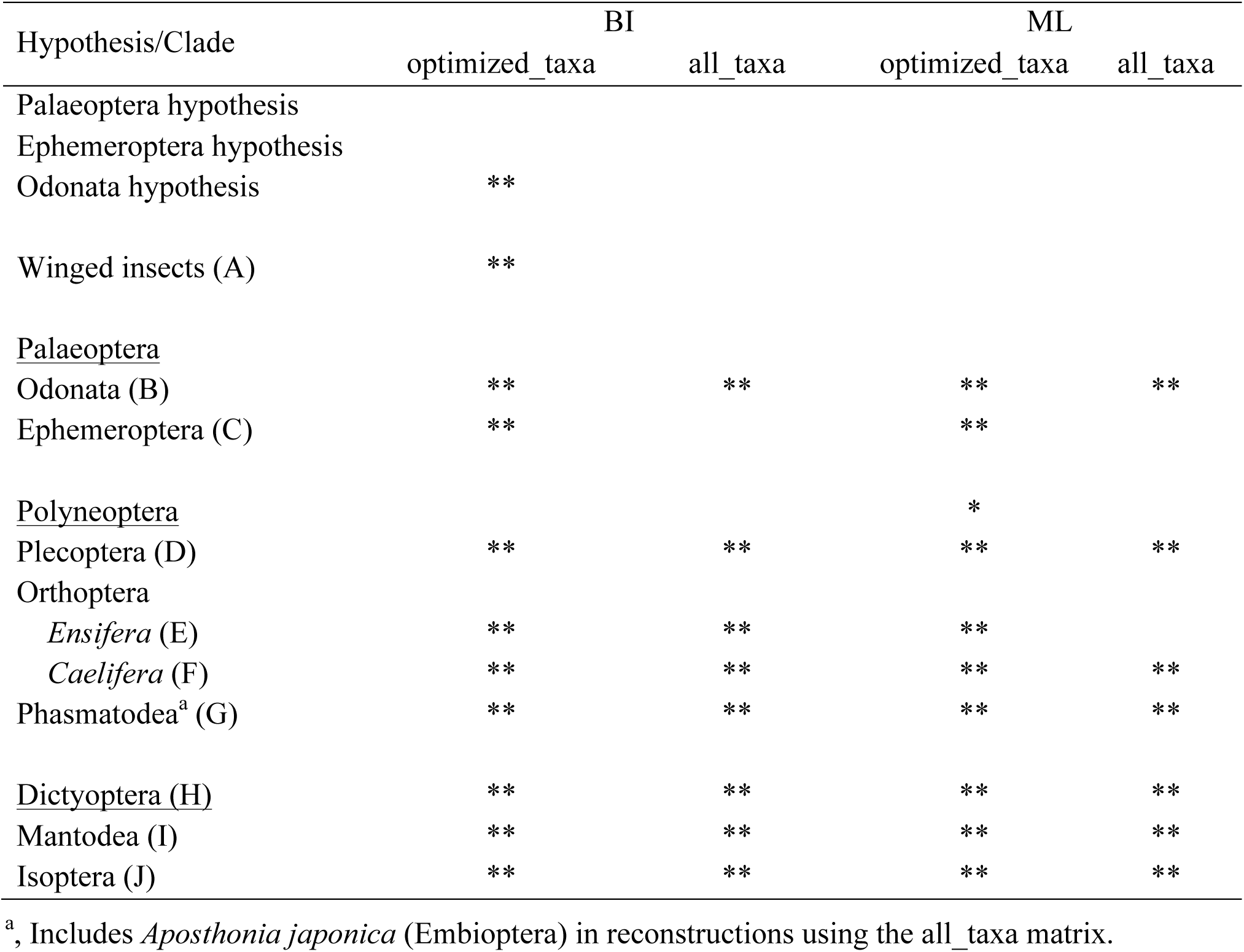
Support for the three hypotheses and node support for clades of interest. Node support is based on Bayesian inference (BI) and maximum likelihood (ML) analyses of two data matrices: optimized_taxa and all_taxa, whereby optimized_taxa had 5 terminal nodes removed based on the occurrence of rogue taxa and long branch attraction. Asterices indicate hypothesis/node support (** = Bayesian posterior probability (BPP) ≥ 0.95 and Bootstrap support (BS) = 90%) * = BPP ≥ 0.90 and BS ≥ 80%). Superorders are underlined, and suborders are italicised. Letters in brackets refer to monophyletic clades in the Fig. 1.

| Hypothesis/Clade | BI |  | ML |  |
| --- | --- | --- | --- | --- |
|  | optimized_taxa | all_taxa | optimized_taxa | all_taxa |
| Palaeoptera hypothesis |  |  |  |  |
| Ephemeroptera hypothesis |  |  |  |  |
| Odonata hypothesis | ** |  |  |  |
| Winged insects (A) | ** |  |  |  |
| <u>Palaeoptera</u> |  |  |  |  |
| Odonata (B) | ** | ** | ** | ** |
| Ephemeroptera (C) | ** |  | ** |  |
| <u>Polyneoptera</u> |  |  | * |  |
| Plecoptera (D) | ** | ** | ** | ** |
| Orthoptera |  |  |  |  |
| *Ensifera* (E) | ** | ** | ** |  |
| *Caelifera* (F) | ** | ** | ** | ** |
| Phasmatodea<sup>a</sup> (G) | ** | ** | ** | ** |
| <u>Dictyoptera</u> (H) | ** | ** | ** | ** |
| Mantodea (I) | ** | ** | ** | ** |
| Isoptera (J) | ** | ** | ** | ** |

<sup>a</sup>, Includes *Aposthonia japonica* (Embioptera) in reconstructions using the all\_taxa matrix.

### 3.2. Phylogenetic reconstruction

The Bayesian reconstruction based on the optimized_taxa matrix produced the tree with most support compared to ML or to either analysis of the all_taxa matrix (Fig. 1, Table 3, Supplementary Figs S1-S3). The basal Odonata hypothesis was highly supported in this tree (Bayesian posterior probability (BPP) = 1.00), whereas neither the basal Ephemeroptera hypothesis nor the Palaeoptera hypothesis received support in any analysis (Table 3). The Odonata were monophyletic in all analyses (BPP = 1.00, BS = 100%) whereas the Ephemeroptera were monophyletic only using the optimized_taxa matrix (Fig. 1, Table 3, Supplementary Fig. S2).

**Fig. 1.**
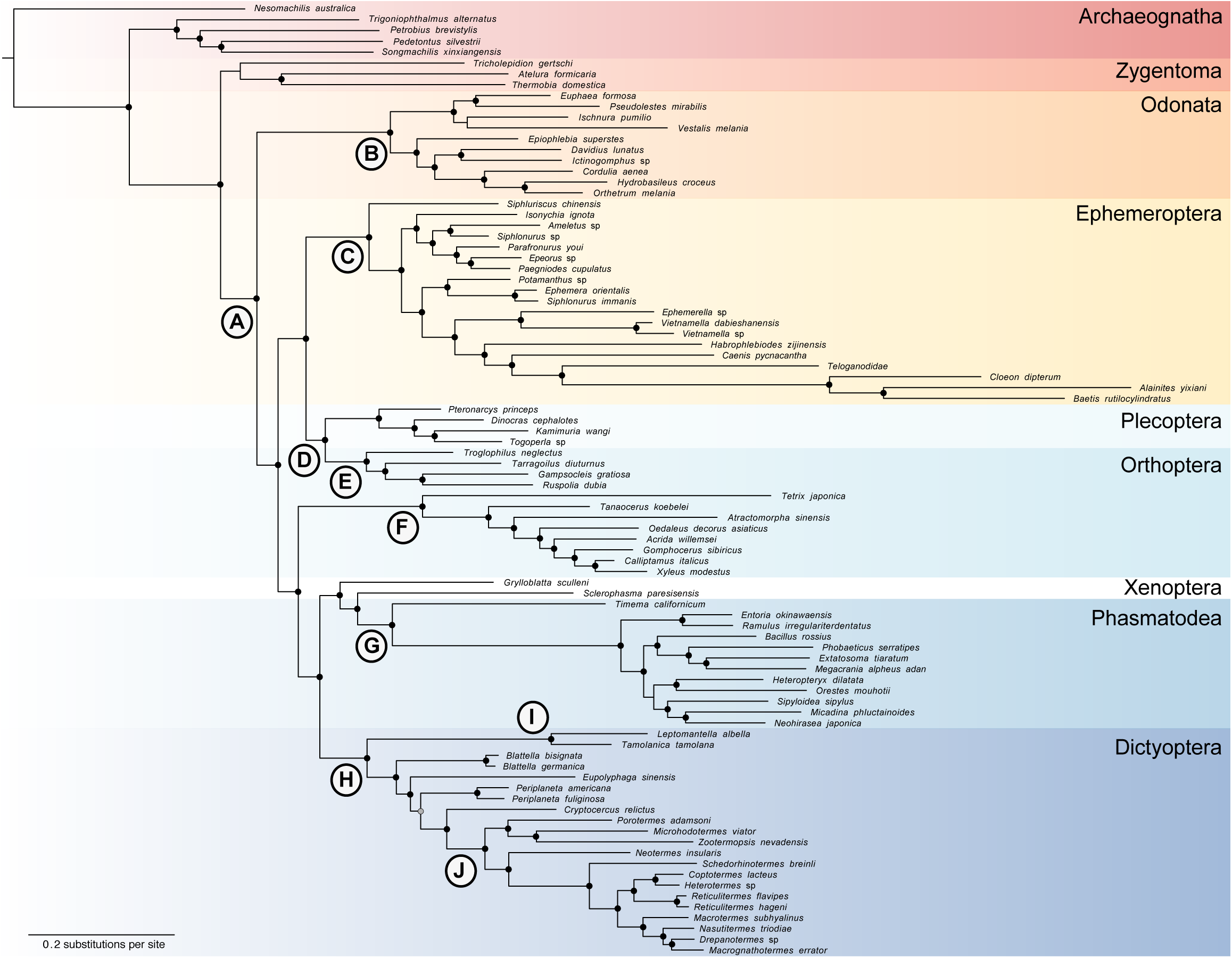
Phylogenetic relationships of insect major orders using Bayesian inference reconstruction based on mitochondrial genome data using the concatenated amino acid sequences of the optimized_taxa matrix (see Table 1). Filled circles indicate supported nodes; whereby black circles represent Bayesian posterior probability (BPP) = 1, and grey circles BPP ≥ 0.95. Letters indicate clades mentioned in the text (see Table 3)

Most other nodes of interest were consistently supported across different matrices and analyses (Table 3). The Polyneoptera were moderately supported (BS ≥ 80%) for the optimized_taxa matrix using the ML. Within the Polyneoptera, all orders except Orthoptera and Blattodea were monophyletic (BPP ≥ 0.95, and BS ≥ 90%, Table 3). The two Orthoptera suborders Ensifera and Caelifera were monophyletic (Table 3), whereby the Ensifera clustered together with the Plecoptera for the optimized_taxa matrix using both ML and BI. The Grylloblattodea, Mantophasmatodea, and Phasmatodea were all well supported using the optimized_taxa matrix. For the all_taxa matrix also the Embioptera were recovered in this clade as sister taxa to the Phasmatodea excluding *Timema californicum* (Fig. 1, Supplementary Figs S1, S3). The superorder Dictyoptera was monophyletic clade, consisting of the monophyletic Mantodea and the Blattodea including the Isoptera.

### 3.3. Mitochondrial genomes

The mitochondrial genome sequences have been deposited at GenBank with the accession numbers XXXXXXXX, XXXXXXXX, and XXXXXXXX. The pyrosequencing run of *C. dipterum* using the 454 GS FLX system resulted in 651,306 reads, of which 1.14% mapped to the assembled mitochondrial genome. The depth of coverage for the *C. dipterum* mitochondrial genome was 249.2x (± 80.6 SD, Fig. 2a). The three mayfly mitochondrial genomes were 15,407 bp (*C. dipterum*), 14,883 bp (*B. rutilocylindratus)*, and 14,355 bp (*H. zijinensis*) long (Table 4). For *B. rutilocylindratus*, the complete mitochondrial genome was sequenced (Fig. 2b). The AT-rich regions were incomplete for genomes of *C. dipterum* and *H. zijinensis*. The three tRNAs between the AT-rich region and *nad2* were also missing in *H. zijinensis* due to the incomplete sequencing (Fig. 2c). All three sequenced mitochondrial genomes contained the entire set of 13 PCGs, two rRNAs, and either 22 tRNAs (*B. rutilocylindratus*, *C. dipterum*) or 19 for the incomplete mitochondrial genome of *H. zijinensis*, including 23 coded at the (+) strand (21 for *H. zijinensis*) and 14 at the (-) strand (13 for *H. zijinensis*) (Fig. 2).

**Table 4.**
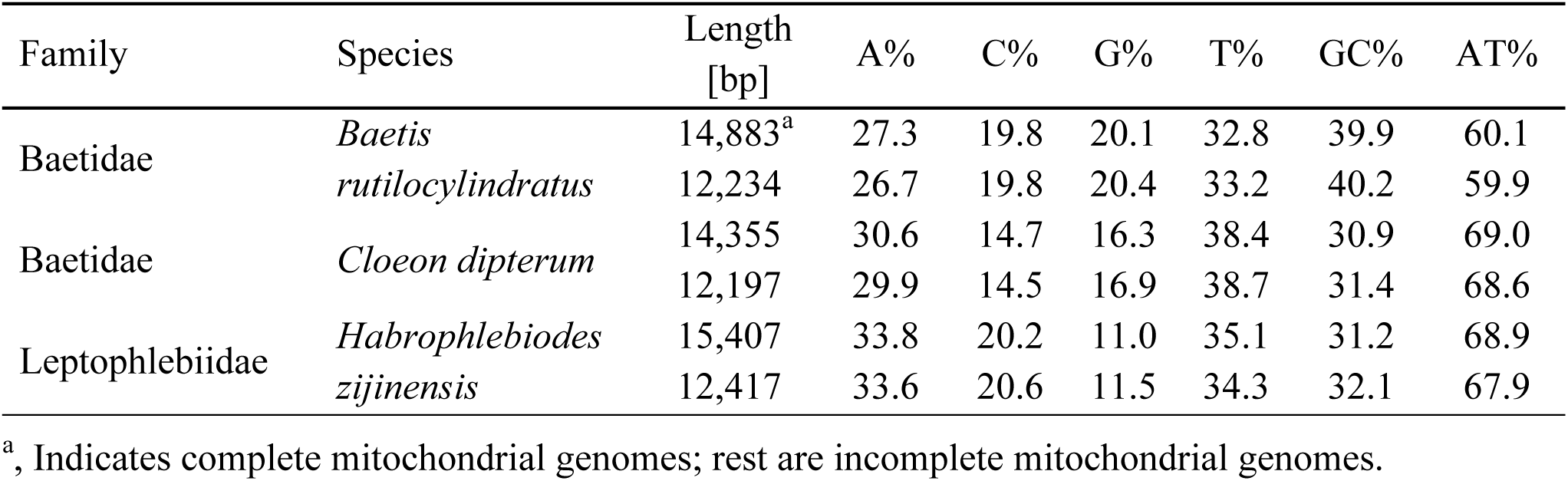
Overview of sequenced ephemeropteran mitochondrial genomes. Total sequence length in base pairs (bp) and individual nucleotide compositions of mayfly mitochondrial genomes calculated based on the whole available sequences and the ones corrected for the incomplete mitochondrial genomes (see 2. Material and methods; for the complete list of all ephemeropteran mitochondrial genomes and Supplementary Table S2).

| Family | Species | Length<br>[bp] | A% | C% | G% | T% | GC% | AT% |
| --- | --- | --- | --- | --- | --- | --- | --- | --- |
| Baetidae | *Baetis* | 14,883<sup>a</sup> | 27.3 | 19.8 | 20.1 | 32.8 | 39.9 | 60.1 |
|  | *rutilocylindratus* | 12,234 | 26.7 | 19.8 | 20.4 | 33.2 | 40.2 | 59.9 |
| Baetidae | *Cloeon dipterum* | 14,355 | 30.6 | 14.7 | 16.3 | 38.4 | 30.9 | 69.0 |
|  |  | 12,197 | 29.9 | 14.5 | 16.9 | 38.7 | 31.4 | 68.6 |
| Leptophlebiidae | *Habrophlebiodes* | 15,407 | 33.8 | 20.2 | 11.0 | 35.1 | 31.2 | 68.9 |
|  | *zijinensis* | 12,417 | 33.6 | 20.6 | 11.5 | 34.3 | 32.1 | 67.9 |

<sup>a</sup>, Indicates complete mitochondrial genomes; rest are incomplete mitochondrial genomes.

**Fig. 2.**
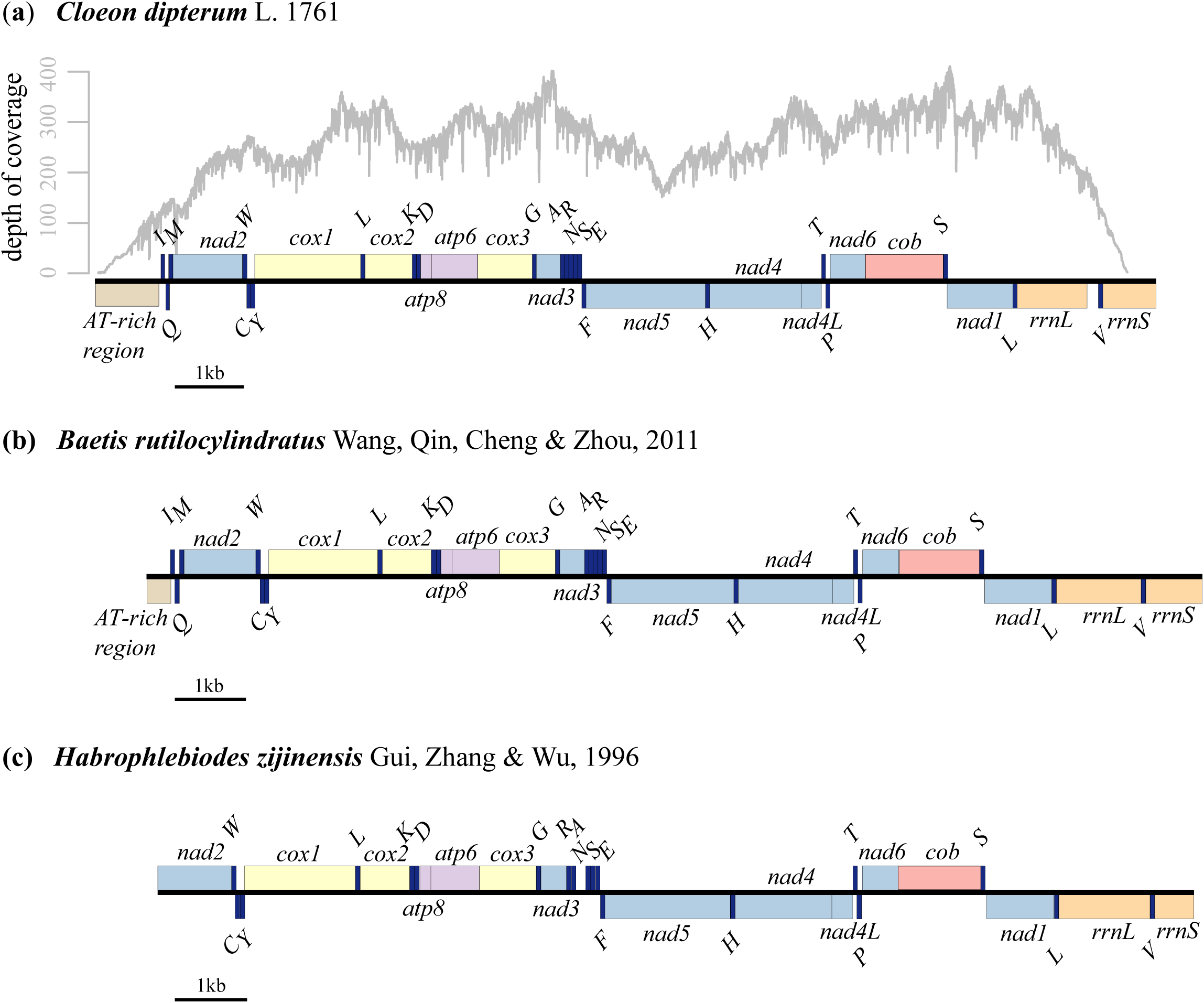
Mitochondrial genome maps of three sequenced Ephemeroptera species. The nearly complete mitochondrial genome of (**a**) *Cloeon dipterum*, including coverage depth, the complete mitochondrial genome of (**b**) *Baetis rutilocylindratus,* and the incomplete mitochondrial genome of (**c**) *Habrophlebiodes zijinensis*. Transfer RNA genes are indicated by single-letter IUPAC-IUB abbreviations for their corresponding amino acid. Protein coding genes and ribosomal RNA genes are listed and colored in the following way: *atp6*, *atp8*, ATP synthase subunits 6 and 8 genes (violet); *cob*, cytochrome oxidase *b* gene (red); *cox1-cox3*, cytochrome oxidase *c* subunit 1-3 genes (yellow); *nad1-6, nad4L*, NADH dehydrogenase subunits 1-6 and 4L (light blue); *rrnS, rrnL*, small and large ribosomal RNA subunits (orange); AT-rich region (brown). Genes located at the (+) strand appear above the central line. Genes located on the (-) strand appear below the central line.

The order of the genes was conserved, with the exceptions of the two inverted tRNAs Arginine and Alanine for *H. zijinensis* (Fig. 2c). All PCGs started with the ATN codons (ATT, ATG, ATA), and mostly ended with the complete termination codon (TAA or TAG). Exceptions were an incomplete T termination codon for *nad4* in *B. rutilocylindratus* and *cox1* started with CTC in *C. dipterum*. The AT content of the mitochondrial genomes that were corrected for the missing regions, was lowest in *B. rutilocylindratus* (59.9%, Table 4, Supplementary Table S2). The mean AT content for all available mayfly mitochondrial genomes was 65.34%. The average AT-skew was −0.03 (± 0.04 SD) and the average GC-skew was −0.13 (± 0.14 SD).

## 4. Discussion

### 4.1. Phylogenetic relationships of Palaeoptera

Our Bayesian reconstruction based on the optimized_taxa matrix provided strong support for the basal Odonata hypothesis. Previous work based on mitochondrial genome data is (partially) congruent with our findings (Lin et al., 2010; Song et al., 2016; Wan et al., 2012). In contrast to our results, a handful of studies based on nucleotide and aa sequences of mitochondrial PCGs and mitochondrial RNA (mtRNAs) found overall strong support for the basal Ephemeroptera hypothesis (Ma et al., 2014; Ma et al., 2015; Song et al., 2016; Zhang et al., 2008). Interestingly, Song et al. (2016), supporting overall the Ephemeroptera hypothesis, found evidence for the Odonata hypothesis when using aa sequence data. Our results confirm the trend towards a better support for the ancestral position of the Odonata when using an increased mayfly taxon sampling (more than ten species per lineage) (Kjer, 2004; Misof et al., 2007; Yoshizawa and Johnson, 2005). The exceptions for this are the work by Ogden and Whiting (2003) and Thomas et al. (2013), that overall support the Ephemeroptera respective Palaeoptera hypothesis but contain some data sets in favor of the Odonata hypothesis. Early inconsistencies based on different sequence alignment strategies (Ogden and Whiting, 2003) appear to become less relevant due to the advances of multiple sequence alignment programs. Overall, the vast majority of studies were based on BI and ML. Thomas et al. (2013) recovered increased resolution using BI in comparison to ML. Different data partitioning (i.e., gene trees resulting in different tree topologies) by Kjer et al. (2006) supported different hypotheses for different genes. Evidently, using rRNAs (nDNA and mtDNA) and the nDNA EF1-a the Odonata hypothesis was recovered whereas the analyses based on the combined and on the mtDNA data matrix resulted in a monophyletic palaeopteran clade. It is questionable whether an increase in analyzed genomic data (i.e., phylogenomic studies) will be able to resolve the Palaeoptera problem. Notably, Misof et al. (2014) and Regier et al. (2010), using 1,478 PCGs and seven palaeopteran species, respective 62 PCGs and four palaeoptera species, recovered the Ephemeroptera and Odonata as sister clades (i.e., the Palaeoptera hypothesis) but without high support values. Overall, more knowledge about the use and limitation of individual markers are needed. For example Simon et al. (2012) found that proteins involved in cellular processes and signaling harbor the most phylogenetic signal.

The increased mayfly taxon sampling highlighted the phylogenetic diversity (as indicated by the branch lengths of the tip taxa, Fig. 1, Supplementary Figs S1-S3) of the Palaeoptera. The overall long-branch lengths and high rate-heterogeneity across sites might be one of the explanations for the “Palaeoptera problem”. Especially the family of the Baetidae, which was long thought to be the most ancestral mayfly family (Ogden and Whiting, 2005), showed high base composition heterogeneity possibly evidencing LBA. However, in agreement with other more recent studies we also recovered *S. chinensis* as sister taxa to all other mayflies (Li et al., 2014; Ogden et al., 2009). It remains to be resolved whether an increased taxon sampling may obviate LBA *sensu* Bergsten (2005).

### 4.2. Phylogenetic relationships of Polyneoptera

Besides the Palaeoptera, a universal consensus on the placement of the Plecoptera, Dermaptera, Embioptera, and Xenoptera (i.e., Grylloblattodea and Mantophasmatodea) remains elusive (Song et al., 2016). The monophyly of the Polyneoptera has become more widely accepted (Ishiwata et al., 2011; Letsch and Simon, 2013; Letsch et al., 2012; Misof et al., 2014; Song et al., 2016; Wan et al., 2012), although several studies and data matrices do not support their monophyly (Kjer et al., 2006; Simon et al., 2012; Song et al., 2016). This is also reflected by our results, recovering the Polyneoptera as moderately supported monophyletic clade when using the optimized_taxa matrix with ML. The placement of the order Plecoptera also remains elusive. The order often clusters among the more ancestral polyneopteran orders (Misof et al., 2014; Song et al., 2016; Wan et al., 2012) and in a close relationship to the Dermaptera, although earlier studies using few species and the aa sequences of the mitochondrial PCGs also recovered a sister relationship to the Ephemeroptera (Lin et al., 2010; Zhang et al., 2009). Notably, Song et al. (2016) was the only study including four dermapteran species. The monophyly of the Orthoptera has been established by previous studies based on mitochondrial genome data and a large number of PCGs (Fenn et al., 2008; Ma et al., 2009; Misof et al., 2014; Song et al., 2016). However, in a recent study, the Ensifera also clustered as more ancestral clade (Song et al., 2016). As for this study, Song et al. (2016) also used BI and ML based on mitochondrial genome data. Notably, Song et al. (2016) explain the more ancestral position of Ensifera by more similar sequence compositional bias in comparison to the outgroups. The relationship between Grylloblattodea and Mantophasmatodea (i.e., Xenoptera superorder) remains elusive. Mostly this is due to the limited amount of available molecular data, including ordinal-level phylogenies based on two (Ishiwata et al., 2011; Song et al., 2016) or three xenopteran species (Misof et al., 2014). Using mitochondrial genome data and one species per order, also Song et al. (2016) did not recover them as a monophyletic clade. Their monophyly was supported by previous studies using one Mantophasmatodea and one Grylloblattodea species (Ishiwata et al., 2011), respective one Mantophasmatodea and two Grylloblattodea species (Misof et al., 2014). The Phasmatodea have also been found in previous studies as paraphyletic clade due to the outside position of *T. californicum* (Ma et al., 2014; Song et al., 2016). In contrast, Misof et al. (2014), using a large set of nuclear PCGs also found *Timema* as the sister species to all other representatives of the order. The monophyly of the superorder Dictyoptera is generally accepted and the hierarchical clustering (Mantodea + (Blattodea + Isoptera)) has been well supported (Inward et al., 2007; Ishiwata et al., 2011; Kjer et al., 2006; Legendre et al., 2015; Ma et al., 2014; Misof et al., 2014; Song et al., 2016; Wan et al., 2012) using nucleotide and aa sequences of both mtDNA and nDNA.

### 4.3. Optimizing data matrix

The optimization of the data matrix overall resulted in a better-supported tree (as measured by the total number of supported nodes and the fact that all nodes supported in the all_taxa matrix were also recovered in the optimized_taxa matrix). In a study on butterfly phylogenetics, the authors also removed rogue taxa and found dramatically increased bootstrap support values (Regier et al., 2013). Comparing the number of supported nodes from the BI and ML approach, the former always resulted in more resolved nodes. This result is consistent with previous work on the “Palaeoptera problem”, recovering more support for analyses using BI (Thomas et al., 2013). Generally, the choice of the outgroup species was reported as being crucial for resolving problematic splits in the tree of life such as insects origin (Thomas et al., 2013). Thus, finding the outgroup as a rogue taxon was perhaps not surprising. The close phylogenetic relationship of the Dermaptera and Ephemeroptera (Supplementary Figs S1, S3) might be misleading due to LBA. Li et al. (2014) also found the order Dermaptera as being closely related to the mayflies. On the other hand studies based on phylogenomic data reported the Dermaptera as sister taxa to the orders Plecoptera (Ishiwata et al., 2011; Ma et al., 2014; Simon et al., 2012; Wan et al., 2012; Wu et al., 2014) Zoraptera (Blanke et al., 2013; Misof et al., 2014). Using mitochondrial genome data and an increased taxon sampling (i.e., four dermapteran species), Song et al. (2016) found the Dermaptera and/or Plecoptera being sister to the remaining taxa within Polyneoptera. Notably, when only using one species of Dermaptera, *C. fletcheri* emerged as sister taxa to the Ephemeroptera (Song et al., 2016). This result in agreement with our findings is presumably due to LBA. Phylogenomic data matrices (e.g., expressed sequence tags data: Letsch et al., 2012, 1,478 PCGs: Misof et al., 2014, and nuclear genes Ishiwata et al., 2011) found the Embioptera to be sister clade to the Phasmatodea. Using mitochondrial genomes data and BI, the Embioptera were also found as sister group to the Phasmatodea (excluding *T. californicum*) (Song et al., 2016). However, in the same study, they also found support for the sister-relationship of Embioptera-Zoraptera using ML (Song et al., 2016). Overall, more phylogenetic studies will be needed to clarify this issue; mostly also because the use of few taxa from anomalous orders (i.e., Dermaptera and Embioptera) tend to evoke LBA (Song et al., 2016).

### 4.4. Characterization of ephemeropteran mitochondrial genomes

Mayfly mitochondrial genomes are widely conserved in gene content and nucleotide composition across eleven families, with only a few differences in the content of their tRNAs. The coverage of the *Cloeon*-mitochondrial genome was in agreement with other studies, using the same sequencing platform (e.g., 59-281x, Pons et al., 2014). The missing of part of the AT-rich region is due to reduced sequencing and assembly efficiency of this low complexity region and common among insect mitochondrial genomes (Li et al., 2014; Tang et al., 2014). The gene order and orientation were with the exception of two shifted tRNAs for *H. zijinensis* identical to the ancestral insect mitochondrial genome (Boore, 1999; Cameron, 2014; Simon et al., 1994). Other mayflies are also reported to miss complete T termination codons in the genes *cox2* and *nad5* (Lee et al., 2009; Li et al., 2014; Tang et al., 2014). The two rRNAs were located between *trnL* and *trnV* (*rrnL*), and between *trnV* and the AT-rich region (*rrnS*), respectively. All ephemeropteran mitochondrial genomes contain an AT-rich region, which is placed between the *rrnS* (- strand) and *trnI* (+ strand). Li et al. (2014) reported two distinct parts within the AT-rich region in *Siphluriscus chinensis,* which also seems to be present in *C. dipterum*. Therein, they described the so called CR_1,_ which is located close to the *rrnS* and has a high AT content (71.6%), including six identical 140 bp sequences, and the CR_2_, which is close to the *trnI* and has a lower AT content (58.1%). *E. orientalis* contains two identical 55 bp long sequences in the AT-rich region (Lee et al., 2009). Few mayfly mitochondrial genomes differ in their gene content from the ancestral insect mitochondrial genome, possessing one additional tRNA. The two heptageniid species *Parafronurus youi* and *Epeorus* sp. encode a second copy of the *trnM* (AUG, *trnM2*) gene located between *trnI* and *trnQ* (Tang et al., 2014; Zhang et al., 2008). For *S. chinensis*, an additional *trnK2* (AAA) gene is described (Li et al., 2014). Overall, Song et al. (2016) found similar genetic distances for the early branching insects lineages. The AT contents for the Ephemeroptera were very similar to the ones of Odonata (62.6%-68.5% for *cox1*, Kim et al., 2014).

## 5. Conclusions

Our analysis based on an increased mayfly taxon sampling and data matrix optimization supported the Odonata hypothesis and unravelled fine structural changes within the palaeopteran mitochondrial genomes. While both BI and ML overall resulted in highly supported trees, only the BI based on an optimized taxa matrix highly supported the sister-relationship of the Ephemeroptera and Neoptera. The optimized taxa matrix, excluding rogue taxa and taxa with LBA, resulted in an overall better supported tree (as measured by the number of supported nodes) for both BI and ML. Our findings highlight the essential need to increase the taxon sampling of under-represented lineages (such as the Palaeoptera) in order to resolve their phylogenetic position. Here we demonstrated the need for an increased taxon sampling in combination with data matrix optimization, and the use of different phylogenetic approaches in order to resolve an ancient radiation. Establishing general recommendation for data matrix optimization requires additional analyses on a broader range of lineages.

## Author contributions

SR and MTM conceived the study; SR sequenced and analyzed the *Cloeon dipterum* genome, and performed all combined analyses; PC and CZ obtained and analyzed the *Baetis rutilocylindratus* and *Habrophlebiodes zijinensis* genomes; SR and MTM wrote the manuscript. All authors made contributions to subsequent revisions and agreed to the final version.

## Acknowledgements

We are grateful to D. H. Funk from the Stround Water Research Center for providing the *Cloeon dipterum* specimens, S. Mbedi, K. Preuß, and L. Wächter for their great help with laboratory work, G. Glöckner, and C. Mazzoni for constructive comments about the analysis of the mitochondrial genomes, and Zedat-HPC at the Freie Universität Berlin, Germany for providing access to high-performance computing clusters. SR thanks the Janggen-Pöhn-Stiftung (http://www.janggen-poehn.ch/) for a Postdoctoral stipend. We also want to thank the members of our research groups for constructive working environments. This work was supported by the Leibniz Association PAKT für Forschung und Innovation (‘FREDIE’ project: SAW-2011-ZFMK-3).

## Data statement

DNA sequences are available on Genbank under accessions no XXXXXXXX-XXXXXXXX. Sequence alignments used for the phylogenetic analyses are available on Mendeley Data https://doi.org/XXX.

